# Cystin is required for maintaining fibrocystin (FPC) levels and safeguarding proteome integrity in mouse renal epithelial cells A mechanistic connection between the kidney defects in *cpk* mice and human ARPKD

**DOI:** 10.1101/2022.04.19.488799

**Authors:** Yiming Zhang, Chaozhe Yang, Wei Wang, Naoe Harafuji, Piotr Stasiak, P. Darwin Bell, Ljuba Caldovic, Elizabeth Sztul, Lisa M. Guay-Woodford, Zsuzsanna Bebok

## Abstract

Autosomal recessive polycystic kidney disease (ARPKD) is caused primarily by mutations in *PKHD1*, encoding fibrocystin (FPC), but *Pkhd1* mutant mice fail to express renal cystic disease. In contrast, the renal lesion in *Cys1*^*cpk/cpk*^ (*cpk*) mice with loss of the cystin protein, closely phenocopy ARPKD. Recent identification of patients with *CYS1*-related ARPKD prompted the investigations described herein. We analyzed cystin and FPC expression in mouse models (*cpk*, rescued-*cpk* (*r*-*cpk*), *Pkhd1* mutants) and cortical collecting duct (CCD) cell lines (wild type (*wt), cpk)*. We found that cystin deficiency led to diminished FPC in both *cpk* kidneys and CCD cells. In *r-cpk* kidneys, FPC increased and siRNA of *Cys1* in *wt* CCD cells reduced FPC. Conversely, FPC deficiency in *Pkhd1* mutants did not affect cystin levels. Cystin deficiency and the associated reduction in FPC levels impacted the architecture of the primary cilium, but not ciliogenesis. Similar *Pkhd1* mRNA levels in *wt, cpk* kidneys and CCD cells suggested posttranslational mechanisms directed FPC loss and studies of cellular protein degradation systems revealed selective autophagy as a possible mechanism. Loss of FPC from the NEDD4 E3 ubiquitin ligase complexes caused reduced polyubiquitination and elevated levels of functional epithelial sodium channel (NEDD4 target) in *cpk* cells. We propose that cystin is necessary to stabilize FPC and loss of cystin leads to rapid FPC degradation. FPC removal from E3-ligase complexes alters the cellular proteome and may contribute to cystogenesis through multiple mechanisms, that include MYC transcriptional regulation.

## INTRODUCTION

The C57BL/6J*Cys1*^*cpk/cpk*^ (*cpk*) mouse is the first described experimental model of polycystic kidney disease (PKD) (1), in which a spontaneous mutation in *Cys1* (encoding the protein cystin) causes renal cystic disease that phenocopies human ARPKD (2). In contrast, causative mutations in the majority of ARPKD patients occur in *PKHD1*, encoding fibrocystin (FPC) (3). Recently, patients with ARPKD linked to mutations in *CYS1* have been identified (4), prompting interest in possible functional relationships between cystin and FPC. It is not understood why mutations in *CYS1 (Cys1)* and *PKHD1* lead to similar kidney phenotypes in humans, but discordant phenotypes when the mouse orthologue, *Pkhd1* is disrupted. While a physical interaction between cystin and FPC has not be demonstrated, in the current study, we sought to examine potential functional linkages between cystin and FPC in *cpk* (4) and *Pkhd1* mutant mouse kidneys (5-7) as well as cortical collecting duct (CCD) cell lines (*wt, cpk*) (8, 9).

The *cpk* phenotype is caused by a frameshift mutation within *Cys1* exon 1, leading to reduced *Cys1* mRNA levels and consequent loss of cystin, a hydrophobic, 145-amino acid, N-terminally myristoylated protein (10). In mouse renal epithelial cells, cystin traffics to the primary cilium and co-localizes with other PKD-associated proteins, including polaris/Ift88 (11). N-myristoylation modulates protein plasma membrane association and impacts protein function via subcellular trafficking and localization (12-16). Cystin is myristoylated at amino acid residue G2 and associates with lipid raft membrane microdomains (17). N-myristoylation facilitates cystin association with the plasma membrane association and a putative myristoyl-electrostatic switch modulates its subcellular trafficking and localization (12-16). Myristoylation and a short peptide sequence between amino acids 28 and 35 (AxEGG) are both required for cystin trafficking to, and retention in, the primary cilium (18). Cystin also localizes to the nucleus in a regulatory complex with necdin (18). In the absence of cystin, necdin activates *Myc* transcription. Importantly, c-Myc levels are significantly elevated in both *PKHD1-*deficient human ARPKD kidneys and mouse *cpk* cystic kidneys (18). Dysregulated *Myc/MYC* expression has been implicated as a central driver of PKD (19, 20).

In contrast to *cpk* mice, *Pkhd1* mutants fail to develop an ARPKD-like renal phenotype (7, 21), for as yet undefined reasons. FPC, encoded by *Pkhd1*, is a large transmembrane protein with an extensive extracellular N-terminal domain, a single pass transmembrane domain and a short cytoplasmic C-terminal domain (CTD) (21, 22). Although FPC function is largely unknown, the FPC-CTD is required for the function of the NEDD4 family of HECT domain-containing E3 ubiquitin ligases and co-localizes in vesicles with the NEDD4 ubiquitin ligase interacting protein, NDFIP2 (23). FPC deficiency is associated with enhanced activity of ENaC (23), a NEDD4 target protein (24), which may contribute to the systemic hypertension that is characteristic of ARPKD (25, 26). However, the influence of FPC on the cellular proteome and the mechanisms through which FPC loss, or reduction alters cellular homeostasis remains unclear. Recent reports have provided new insights into the formation of E3 ligase complexes, determinants of substrate specificity, and regulation of the cellular proteome through ubiquitination (27-29). Based on these developments, we speculate that FPC, through its function in E3 ligase complexes, plays a crucial role in maintaining the cellular proteome and disruption of proteome homeostasis may contribute to ARPKD renal pathogenesis.

In the current study, we examined cystin and FPC levels in the kidneys of *wt, Pkhd1* mutant (7, 21) *cpk* and rescued *cpk* (r-*cpk*) mice with collecting duct specific expression of a wild-type, green fluorescent protein (GFP)-tagged *Cys1* transgene, which reduced cyst formation in our previous studies (4). FPC was markedly reduced in *cpk* compared to *wt* kidneys and FPC abundance increased in r-*cpk* kidneys. Conversely, FPC loss did not alter cystin levels in *Pkhd1* mutants. Using previously developed and characterized renal CCD cell lines generated from *wt* and *cpk* mouse kidney CCD cells (8, 9), we observed that FPC levels were reduced in cystin deficient-cells. FPC loss led to a significant, generalized reduction of ubiquitination that is likely to alter cellular proteome. As would be expected based on previous reports (23), ENaC levels were elevated and sodium transport was increased in *cpk* cells, consistent with inhibited degradation of ENaC, a NEDD4 target protein (30).

Based on our observations, we propose that cystin is necessary to stabilize FPC and regulate the cellular proteome. Further, we propose that the ARPKD-like renal cystic phenotype in *cpk* mice results from the combined loss of cystin and FPC, leading to proteome alterations that disrupt cellular homeostasis. In addition, the loss of cystin-dependent negative regulation of *Myc* transcription results in overexpression of Myc that can drive cyst formation (18). While the current studies were not designed to directly address the discordant renal phenotypes observed in *PKHD1*-associated human ARPKD and mouse *Pkhd1* mutants, identification of a functional connection between cystin and FPC and demonstration that FPC loss does not reduce cystin levels leads us to speculate that in *Pkhd1* mutant mice, but not in human *PKHD1*-deficient kidneys, cystin prevents the development of the ARPKD-like renal cystic phenotype because cystin acts to constrain *Myc* transcription (18, 31).

## MATERIALS AND METHODS

### Mice

Kidney tissues were collected from 14-days old, male animals. Animal procedures were approved by the Institutional Animal Care Committee of Children’s National Research Institute.

### Analysis of kidney tissue lysates

Kidney tissues were collected, homogenized, and processed for immunoblotting as previously described (18). Immuno-reactive protein bands were visualized using SuperSignal West Dura chemiluminescent substrate (Thermo Fisher Scientific # 34076) and images were obtained using a ChemiDoc Imaging System (Bio-Rad laboratory, Inc.). Densitometry data were collected using Image Lab (Bio-Rad laboratory, Inc., Version 6.0) and statistical analysis was performed using Graph Pad Prism software.

### Cell lines

mTERT-immortalized mouse CCD cells were cultured as described previously (8, 9) in 1:1 DMEM/F-12 (Gibco GlutaMAX™; Fisher #10565018) with 5% FBS (Gibco One Shot™; Fisher #A3160602) and Penicillin/Streptomycin (Mediatech, Inc., Corning 30-002). Complete cell culture medium was supplemented with dexamethasone (0.2 μg/ml; Sigma #D8893-1mg), triiodothrionine (10 nM; Sigma #T5616-1mg), 1x insulin-transferrin-sodium selenite (Sigma #I1884-1VL), and L-Glutamine (200mM; Corning Cellgro #25-005-CI).

### Real-time PCR analysis

Relative quantification of *Pkhd1* mRNA in cells was performed using the delta-T method (32) using the following primer sets from Integrated DNA Technologies: *Pkhd1*-T forward 5’-CAGTTCTTGCCAGAGCATTTAC-3’, *Pkhd1*-T reverse 5’-CAGAATCTCACCTCCTGCTATG-3’, *Pkhd1*-M forward 5’-CTGATAATGCACAGGGACCTAC-3’, *Pkhd1*-M reverse 5’-GGCAAAGGATGAAATGGAAGT G-3’, *Pkhd1*-B forward 5’-CCA CCAGAAACCATCCAGTAA-3’, *Pkhd1*-B reverse 5’-AGACCTCTCCTCTCCCATTT-3’. Relative *Pkhd1* mRNA levels from kidneys were determined by measuring *Pkhd1* ex66-67 expression normalized to *Ppia* (*peptidylpropyl isomerase A*); data are expressed as mean ± S.E; n=3 per group, using primers *Pkhd1* forward 5’-CCAGAAGACATATCTGAATCCCAGGC-3’ (mPkhd1 e66F), reverse 5’-AGCAAGAGATCCTGGAACACAGGT-3’ (mPkhd1 e67R), *Ppia* forward 5’-AGCACTGGAGAGAAAGGATT, reverse 5’-ATTATGGCGTGTAAAGTCACCA.

### Antibodies

Primary antibodies used for WB analysis were produced against: FPC, (rat monoclonal (mAb), Baltimore PKD Center; PD1E1; 1:1000), cystin (rabbit, polyclonal (pAb), 70053; 1:1000) (17, 18), PKD2 (rabbit mAb, Baltimore PKD Center; 3374; 1:1000), β-actin (mouse, mAb, Santa Cruz; sc-47778; 1:3000), Ift88 (rabbit pAb, GN593; 1:500) (33), GAPDH (rabbit pAb, Abcam; ab9485; 1:1000), p62/SQSTM1 (mouse mAb, R&D Systems; MAB8028; 1:1000), pan-polyubiquitin (rabbit pAb, Biomol: UG9510; 1:1000), polyubiquitin K63 (rabbit mAb, Millipore; 05-1308; 1:1000), polyubiquitin K48 (rabbit pAb, Cell Signaling; 4289; 1:1000), ENaCα (rabbit pAb, StressMarq; SPC-403; 1:1000). ENaCα (goat pAb, Santa Cruz; sc-22239; 1:200), LAMP1 (mouse mAb: Sztul Lab; 1:20 (34)).

### Western blotting

Cells were lysed in RIPA buffer (Pierce; Thermo #89901) containing protease and phosphatase inhibitors (Halt; Thermo #1861281), sonicated, and centrifuged at 12000 rpm for 30 min. The supernatant was collected, and protein concentration measured using a BCA Protein Assay (Pierce; Thermo # 23225). Equal amounts of proteins were denatured using NuPAGE reducing agent (Invitrogen; Thermo #NP0009), separated by SDS-PAGE using NuPAGE 4-12% bis-tris gel (Invitrogen; Thermo #WG1402BOX) or 3-8% tris-acetate gel (Invitrogen; Thermo #WG1602BOX), and electro-transferred to a nitrocellulose membrane (Odyssey; Li-Cor #926-31092). Membranes were incubated with primary antibodies overnight at 4 0C and secondary antibodies (Li-Cor; 1:20000) for 1 hour at room temperature. Results were imaged using a Li-Cor Odyssey CLx imaging system.

### Cell count

DAPI/Hoechst containing layer was selected and processed. Histogram equalization was performed to enhance the contrast between nuclei and the background (35). After creating a binary mask of the nuclei, the periphery of the objects in the binary mask were dilated (36), and small holes inside the objects were filled (37). The edges of the nuclei were detected and traced using the *bwperim* function (38). Lastly, using the *bwboundaries* function, the number of objects in the binary mask was counted and recorded as the number of cells in the image (36).

### Cilia count

Cilia development was induced by serum starvation for 8 hours as described previously (39). Acetylated α tubulin (AT) staining was used to identify cilia and the percentage of cells with AT-stained primary cilia was plotted relative to total cell number in a viewing area.

### Cilia length and thickness

These parameters were determined using ImageJ. AT-labeled cilium length and thickness was measured in pixels and converted into µm. Results of measurements were plotted and analyzed in GraphPad Prism.

### Densitometry and statistical analysis

Image processing and densitometry were performed using Image Studio Lite software. Results were transferred to GraphPad Prism and data subjected to Student’s t-test or ANOVA analysis.

### Immunocytochemistry and confocal microscopy

Cells were seeded on glass coverslips pretreated with HistoGrip (Fisher #008050) to enhance cell adhesion. After reaching approx. 60% confluence, the cells were washed with PBS, fixed with 4% paraformaldehyde, quenched with ammonium chloride, permeabilized with 0.1% Triton X-100, blocked with 2.5% goat serum, incubated with primary antibodies and Alexa Fluor-conjugated secondary antibodies (1:500), stained with Hoechst 33258 or DAPI, and mounted onto glass microscope slides. Following each step, the cells were washed with PBS. Imaging was done using a Nikon A1R Confocal Microscope and Advanced NIS-Elements software.

### Patch Clamp

Whole cell voltage-clamp recordings were performed on *wt* and *cpk* cells at 24 to 32 hours after seeding on plastic coverslips in regular gro*wt*h medium as previously described (40, 41). Briefly, cells were bathed in an external solution containing (in mM): 145 NaCl, 2.7 KCl, 1.8 CaCl2, 2 MgCl2, 5.5 glucose, and 10 HEPES, pH 7.4 and continuously perfused (∼3 ml/min). Patch pipette resistance was 2.5-3 MΩ when filled with an internal solution containing (in mM): 135 potassium glutamic acid, 10 KCl, 6 NaCl, 1 Mg2ATP, 5.5 glucose, 10 HEPES and 0.5 EGTA, pH 7.3. Junction potentials were considered and corrected before breaking the cell membrane and the formation of a whole-cell patch. To acquire individual cell capacitance, a transient current was induced by applying a short (15 ms) depolarization pulse from -30 to -20 mV before whole-cell current recording. To measure amiloride sensitive currents in voltage clamp, 10 µM amiloride was applied after recording control current. Whole-cell currents were evoked using a voltage step protocol from -90 to 30 mV in 20 mV increments of 450 ms duration at 5 s intervals. The cell membrane potential was held at -30 mV during recording. Signals from whole-cell recording were filtered and sampled at 2 kHz. Current voltage (I-V) relationships were constructed by measuring the steady-state current values at the end of each voltage step. Data acquisition and analysis were performed using Axonpatch 200B amplifier, Digidata 1322A analog-to-digital converter, pClamp 9.2 software (Axon Instruments/Molecular Devices, USA) and Origin Software (Microcal Software, Northampton, MA). All experiments were performed at room temperature.

## RESULTS

### I. Lack of cystin leads to reduced FPC levels in *cpk* kidneys and CCD cells Cystin deficiency is associated with marked reduction of FPC in *cpk* mouse kidneys

We hypothesized that similar *CYS1*/*Cys1* and *PKHD1* renal disease phenotypes reflect disruption of an essential functional interplay between cystin and FPC. In the current study, we compared FPC and cystin levels in the kidneys of *wt, cpk* and r-*cpk* mice, a partially rescued cystic phenotype with collecting duct-specific expression of the *Cys1-GFP* transgene (4) ***(Fig.1A)***. Our results showed a marked (∼90%) reduction ***(Fig.1B)*** of FPC in *cpk* kidneys compared to *wt*. We observed a 3-4-fold increase in FPC levels in r-*cpk* kidneys. Partial rescue of FPC expression in *r-cpk* kidneys is consistent with diminished, but not completely eliminated cyst formation in *r-cpk* mice, which is largely confined to proximal tubular segments (4). In addition, the residual renal cystic disease may also result from diminished function of the cystin-GFP fusion protein (42, 43). Endogenous cystin was present in *wt* but not in *cpk* and *r-cpk* kidneys ***(Fig.1A, lower panel)***. Measurement of *Pkhd1* mRNA levels indicated ∼1.5-fold increase in *cpk* kidney lysates compared to *wt*. This increase, rather than reduction in *Pkhd1* mRNA implies that FPC downregulation in *cpk* mice was not the result of reduced transcription and/or mRNA stability, but rather occurred at the protein level ***(Fig.1C)***.

**Fig.1.**
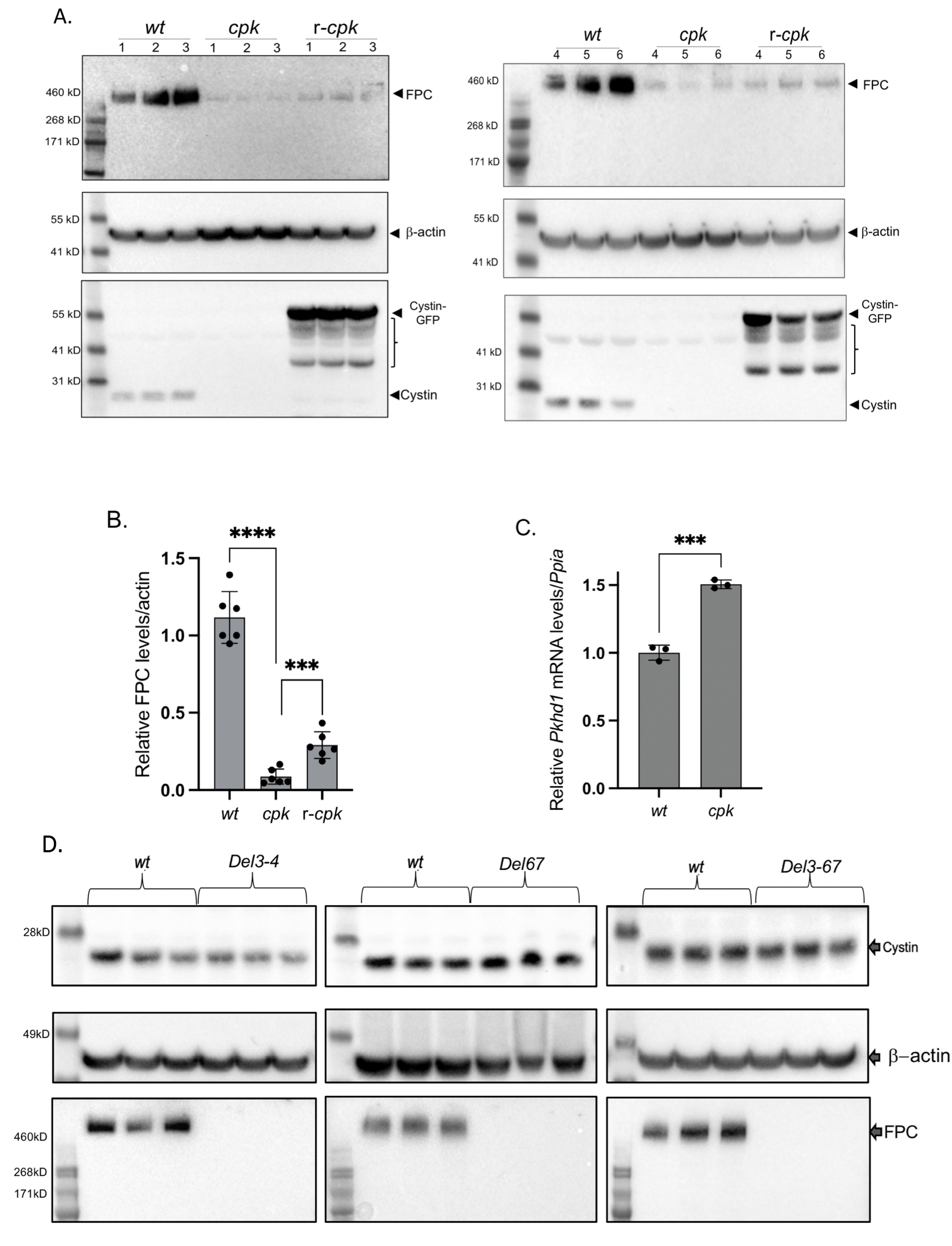
*Cystin deficiency in cpk kidneys leads to FPC loss, with partial rescue of FPC following cystin-GFP expression (r-cpk), but FPC loss (Del3-4, Del67* and *Del3-67) does not alter cystin expression in mouse kidneys*. **A: WB experiments demonstrate significantly lower FPC levels in *cpk, compared to wt cells*, with partial rescue of FPC levels in r-*cpk* kidneys**. Whole kidney lysates (N=2, n=6, male mice) from *wt* (C57BL/6), *cpk* and r-*cpk* mouse kidneys were analyzed for FPC (anti-FPC, C-terminal, rat monoclonal antibody, top), β–actin (loading control, middle) and cystin (anti-cystin polyclonal, rabbit antibody, bottom). In *wt* samples cystin is present (arrow). In *r-cpk* samples a cystin-GFP fusion protein (cystin-GFP, arrow) and multiple unspecific bands are also present (brackets). No endogenous cystin is present in *cpk* and *r-cpk* kidneys. **B: Relative expression of FPC in *wt, cpk* and *r-cpk* kidneys**. FPC levels are plotted relative to β-actin (****: n=6, p<0.0001, ***: p<0.0005, unpaired *t*-test). **C: Higher relative *Pkhd1* mRNA levels in *cpk* kidneys suggest FPC loss at the protein level**. *Pkhd1* mRNA levels were measured by real time RT-PCR. Relative levels of *Pkhd1* mRNA were higher in *cpk* kidneys than *wt* (n=6 ***; p<0.001, unpaired *t* test). **D: Cystin levels are not reduced in *Pkhd1* mutant (*Del3-4, Del67* and *Del3-67)* kidneys**. Whole kidney lysates from *Pkhd1* mutant kidneys (*Del3-4, Del67 and Del3-67*) were tested for cystin and FPC expression and compared to age and gender matched wild-type *(wt)* kidney lysates (n=3, male mice). Cystin expression was similar *wt* and mutant (*Del3-4, Del67 and Del3-67*) kidney lysates (top). β–actin was analyzed as loading control (middle). FPC levels were compared in the same lysates (bottom). Loss of full-length FPC was observed in *Del3-4, Del67 and Del3-67* kidneys.

Because FPC was severely reduced in cystin-deficient kidneys, we tested whether cystin levels were dependent on FPC expression. We analyzed *Pkhd1* mutant (*del3-4, del67, del3-67*) whole kidney lysates for cystin and FPC expression and found that in these *Pkhd1* mutant models loss of full length FPC did not affect cystin expression ***(Fig.1D)***. Taken together, our results demonstrate significant FPC reduction in cystin-deficient kidneys and restoration of FPC levels following expression of a cystin-GFP fusion protein in r-*cpk* kidneys. Conversely, loss of full-length FPC in *Pkhd1* mutant kidneys did not cause reduction in cystin abundance.

#### FPC levels are linked to cystin expression in cortical collecting duct (CCD) cell lines

In subsequent experiments, we examined cell lines generated from *wt* and *cpk* mouse CCD cells (8, 9) to better define the functional connections between cystin and FPC. We validated the nephron-origin of these cell lines by assessing expression of the epithelial cell marker, e-cadherin, and the CCD marker, AQP2 (44) in multiple samples obtained from cell cultures of different passage numbers. We found similar levels of e-cadherin and AQP2 in *wt* and *cpk* cells ***(Fig.2A***,***B)***. As expected, cystin was present in *wt*, but absent from *cpk* cells ***(Fig.2C)***. Similar to our observation in whole kidneys, FPC levels were greatly reduced (by 83±6 %) in *cpk* cells ***(Fig.2D)***. To determine whether the impact of cystin deficiency was FPC-specific or extended to other ciliary and PKD-associated proteins, we examined levels of polycystin-2 (Pc2) and Ift88, proteins previously shown to colocalize with cystin in the primary cilium (10). Reduction of Pc2 abundance (30±6 %) in *cpk* cells was less severe than FPC ***(Fig.2D)***, consistent with previous reports that demonstrated Pc2 reduction in *Pkhd1* knockout kidneys (45). However, lack of cystin did not affect Ift88 levels ***(Fig.2E)***. These results are consistent with our observations in *cpk* kidneys (***Fig.1***) and support a specific relationship between cystin and FPC expression in CCD cells. *Pkhd1* mRNA levels in *wt* and *cpk* CCD cells were similar (***Fig.2F)*** and support the hypothesis that loss of cystin affects FPC abundance post-translationally.

**Fig.2.**
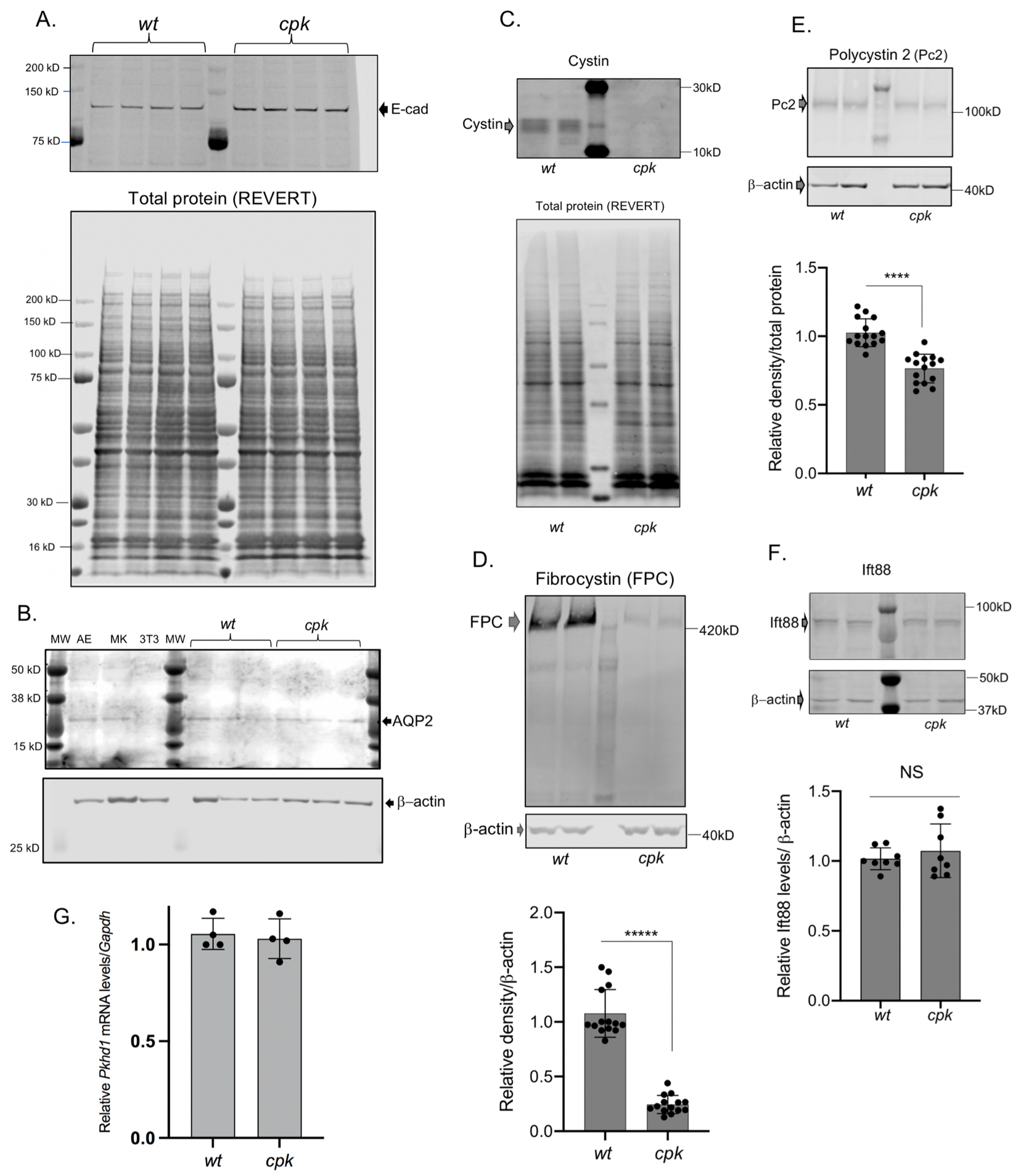
Reduced FPC levels in cpk, compared to wt CCD cells. ***A-B: Characterization of wt and cpk CCD cell lines*** (n=4). **A:** E-cadherin (E-cad) expression was tested as epithelial marker. Cell lysates were randomly selected (*wt, cpk*) from experiments performed in a 6-month period (n=4). **B:** Aquaporin-2 (AQP2) expression (CCD marker, top) in randomly selected from *wt* and *cpk* cell lysates. AE: airway epithelial cells (positive control), MK: mixed mouse kidney epithelial cells, 3T3: mouse fibroblast cell line (negative control). β–actin as loading control (bottom). ***C: Cpk cells lack cystin***. Cystin is only present in *wt* cells. Cystin was labeled with rabbit polyclonal anti-cystin antibody (top). Total protein stain (REVERT) is presented in the lower panel to demonstrate equal loading. ***D: Reduced FPC levels in cpk cells compared to wt***. 30 ug of total protein analyzed by WB using an anti-FPC C-terminal, rat monoclonal antibody (representative gel on top). Relative FPC abundance was quantitatively assessed by densitometry and expressed relative to β-actin (bottom). Experiments were performed using cell culture lysates from different passage numbers (N=7, n=14, unpaired *t* test). ***E: Pc2 reduction in cpk cells***. Pc2 WB using a rabbit polyclonal Pc2 antiserum (top). Pc2 abundance was quantitatively assessed by densitometry and expressed relative to β-actin. Same cell lysates as for FPC were tested for Pc2 (N=7, n=14, ****: p<0.0001, unpaired *t* test). ***F: Ift88 levels were similar in wt and cpk cells***. Ift88 was detected using anti-Ift88 rabbit polyclonal antiserum. No significant differences were observed based on quantitative assessment of densitometry expressed relative to β-actin (N=4, n=8, NS, unpaired *t* test). Same lysates were tested as for Pc2. ***G: Similar relative Pkhd1 mRNA levels in wt and cpk cells***. Plotted values indicate mRNA levels in four independently-derived samples (from cell cultures at different passage number, n=4, NS, unpaired *t* test).

We showed that over-expression of cystin-GFP in *r-cpk* kidneys increased FPC levels (***Fig.1A-B)***. To confirm a role for cystin in maintaining FPC levels and to eliminate possible off-target effects of transient cystin overexpression on cellular functions, we used siRNA to reduce *Cys1* expression in *wt* cells and evaluated both cystin and FPC levels. We achieved siRNA dose-dependent (20-80 ng/ml) reductions in cystin levels 72 hours after siRNA transfection. Reduction of FPC correlated with the efficiency of *Cys1* knockdown (***Fig 3A,B***). Taken together, our *in vivo* and *in vitro* observations strongly support a role for cystin in maintaining FPC levels in both mouse kidneys and renal epithelial cells.

**Fig.3.**
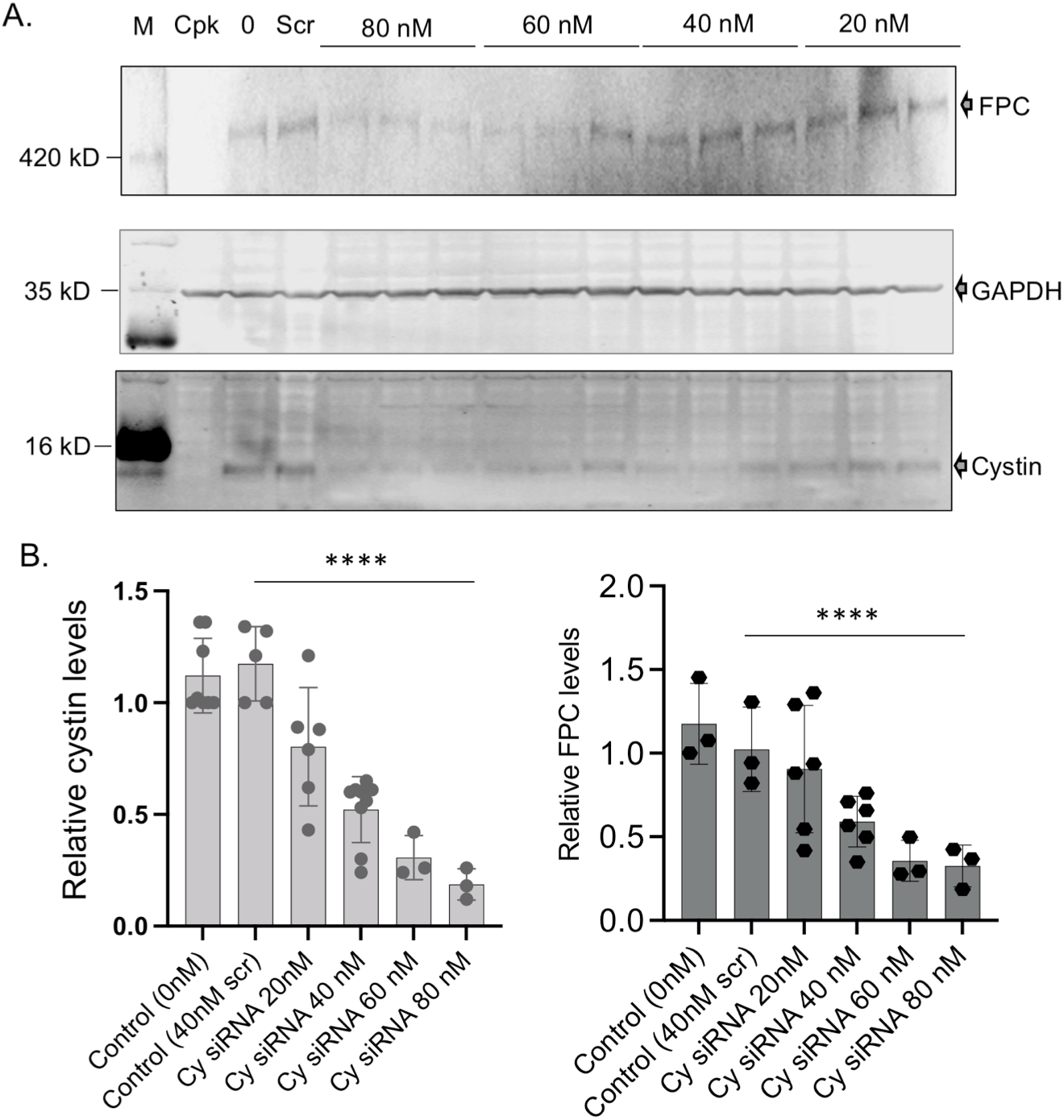
siRNA-based cystin depletion leads to FPC reduction. **A:** Representative WB of FPC (top) and cystin (bottom) following cystin siRNA treatment for 72 hours (siRNA concentrations indicated above image). GAPDH levels (middle) were used to normalize cystin and FPC expression. Cystin expression was measured using anti-cystin rabbit polyclonal antibody (bottom). **B: Efficiency of cystin depletion (left) and FPC levels in the same samples (right)**. Values are plotted relative to GAPDH from n=1-3 separate experiments, with n=3-6 under each siRNA concentration (****p<0.0001, unpaired *t* test, significance was determined between scrambled and cystin-specific siRNA (80 nM)).

#### Cystin and FPC do not colocalize in *wt* cells

Cystin has been localized to the primary cilium in *w*t kidneys (11) and to the nucleus following overexpression (18). However, comparative intracellular localization of endogenous cystin and FPC has not been analyzed in mouse CCD cells (*wt* and *cpk*). Therefore, we performed immunocytochemistry using the same antibodies as in the biochemical studies ***(Figs 1-3)***, co-labeling cystin (green) with FPC (red) ***(Fig.4A)*** in the same cells. In *wt* cells, cystin localized to the cytoplasm and the nucleus in punctuate patterns, whereas FPC labeling showed a vesicular pattern in the cytoplasm. There was no colocalization of the two proteins. Using identical experimental conditions for *cpk* cells, we did not observe cystin labeling and only minimal FPC staining in a small fraction of cells. These results are consistent with the total loss of cystin and significant reduction of FPC in *cpk* kidney-derived cells.

**Fig.4.**
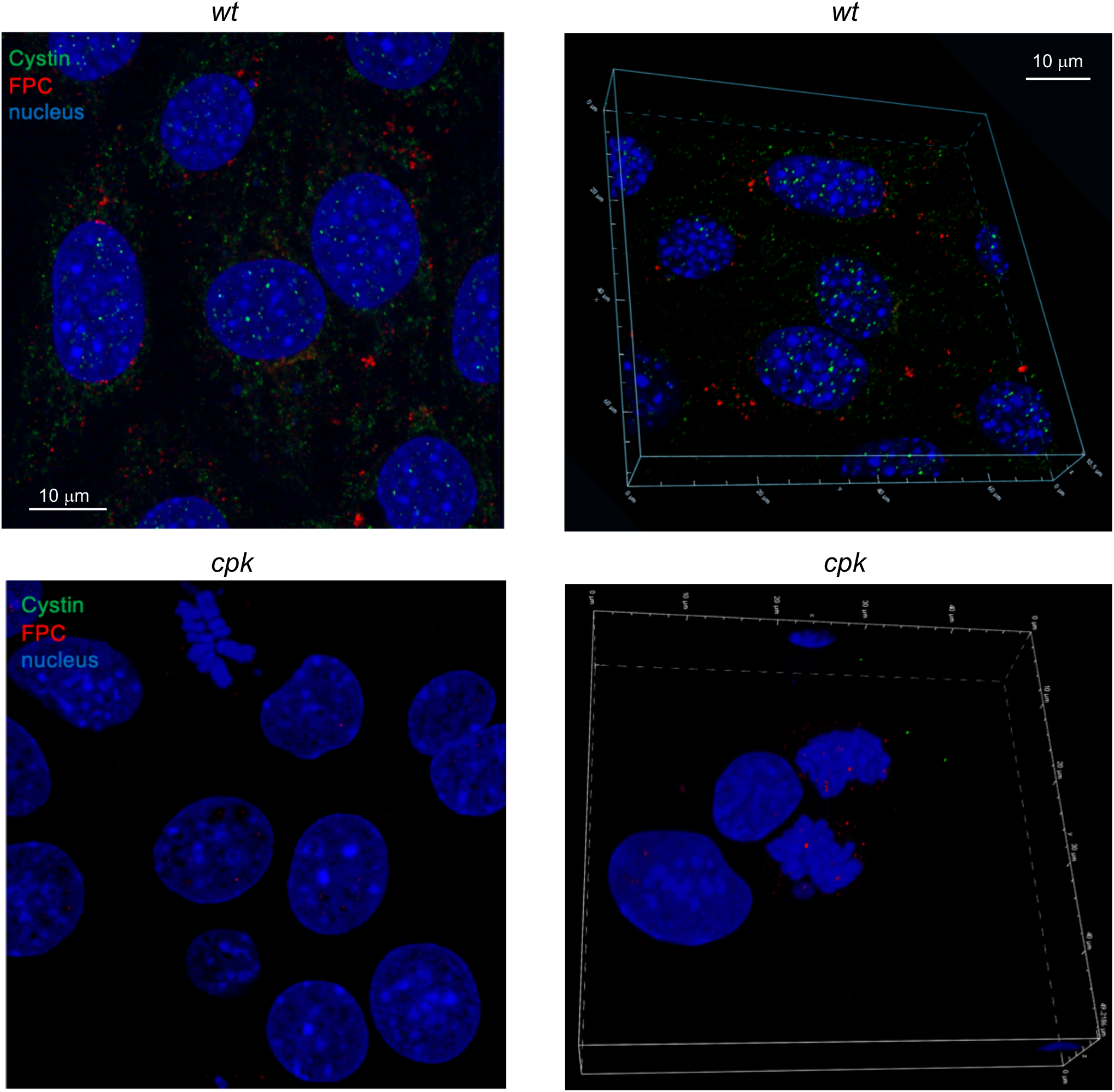
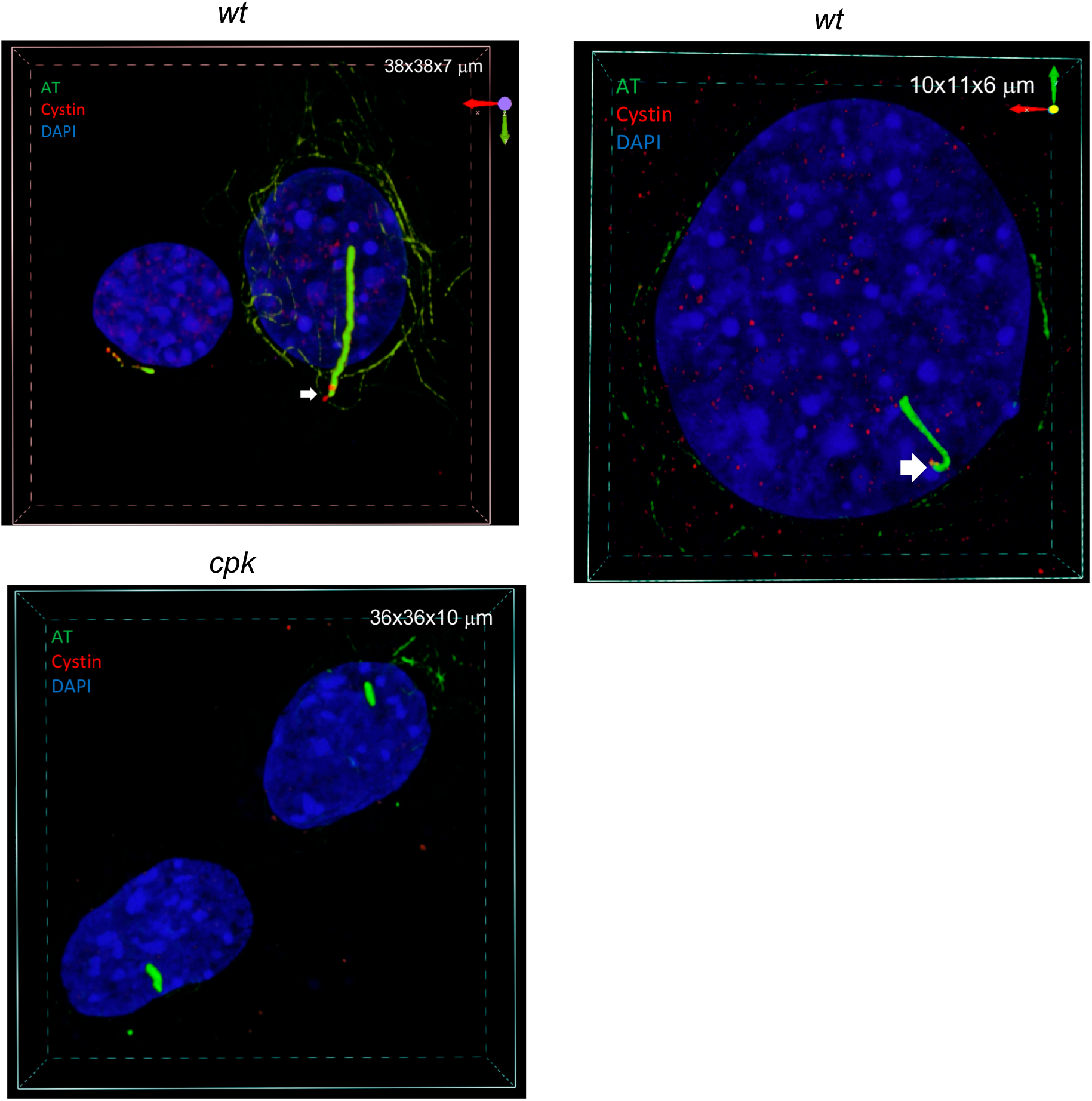
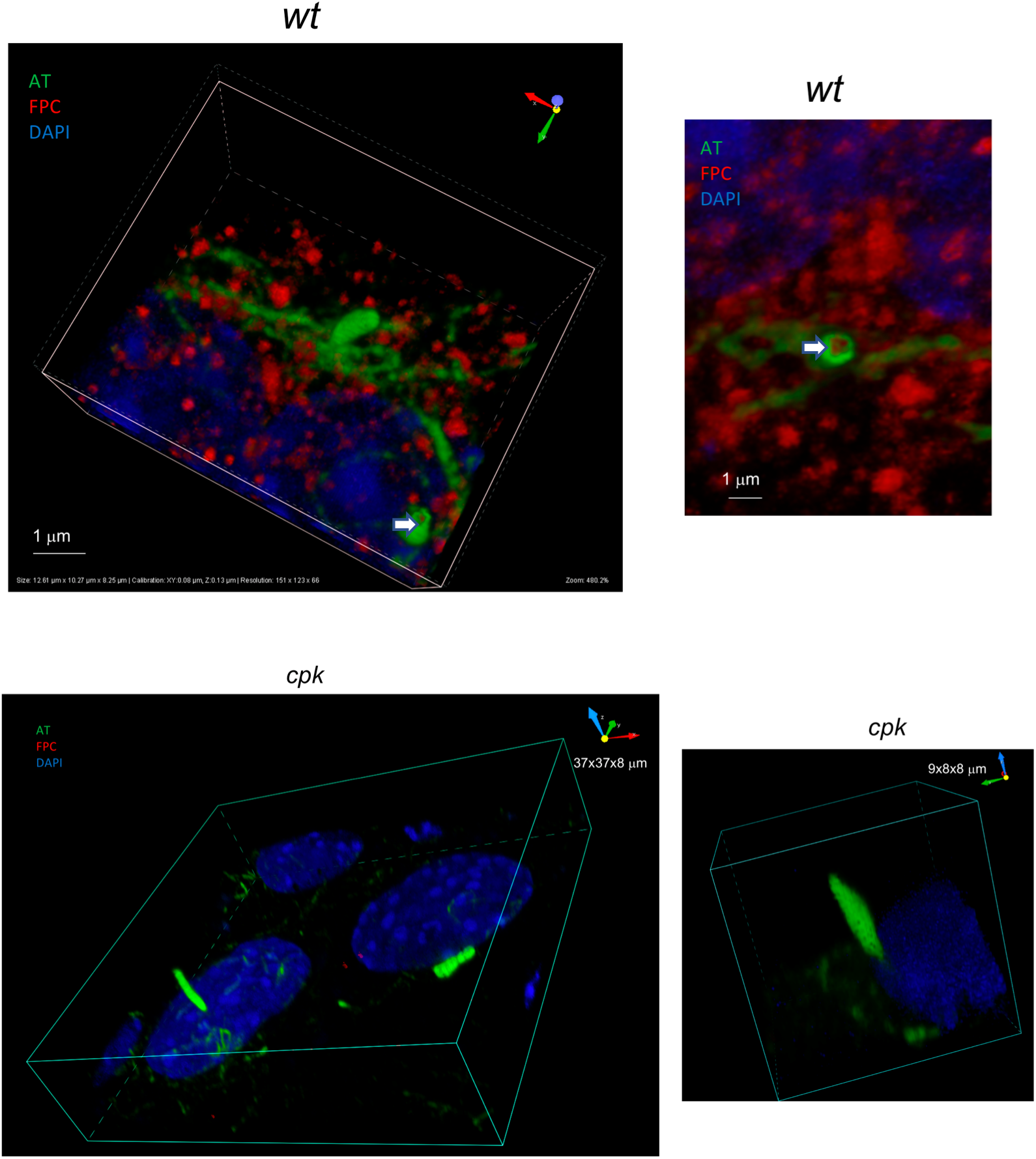
Confocal microscopy of cystin and FPC in wt and cpk cells. ***A:* Cystin (green) and FPC (red) co-staining**. Cystin and FPC do not colocalize in *wt* CCD cells. Representative images of *wt (top)* and *cpk* (bottom) cells and 3D reconstituted images are shown (right). Note cystin in the cytoplasm and nuclei of *wt* cells. No cystin labeling in cpk cells. FPC labeling shows a vesicular pattern in *wt* cells (top) and only minimal FPC labeling in *cpk* cells (bottom). ***B. Co-labeling of cystin (red) with the cilary marker acetylated tubulin (AT, green)***. Images concentrate on the cilia and the nuclei of the cells. Cystin is present in the cilia and the nucleus in *wt* cells. *Cpk* cells lack cystin, but show cilia formation (AT, green). Cystin (red) is localized to the cytoplasm, nucleus and the base of the cilia in *wt* cells (top). Two different cells are shown. Higher cystin levels were observed both in the cytoplasm and in the nucleus in the cell on the right. In cpk cells, no cystin staining was observed, but cilia were present. 3D reconstituted images. ***C*. FPC is present in the cilia of *wt* cells**. Confocal images of WT cells with AT (green) and FPC (red). Optical sectioning of the cilia shows FPC staining in the center of the cilium (arrows). *Cpk* have only minimal FPC labeling in a small fraction of cells.

To examine cystin and FPC localization in the primary cilium, we induced cilia formation (39) and co-labeled either cystin or FPC with acetylated tubulin (AT, a ciliary marker) (46). In *wt* cells, cystin localized to the cytoplasm, the cilia ***(Fig.4A, left)*** and within the nucleus, with variable intensity among the cells (***Fig.4B top***). *Cpk* cells lacked cystin staining in all compartments, but cilia were present (shown by AT labeling) ***(Fig.4B, bottom)***. FPC labeling showed similar pattern in ciliated cells (***Fig.4A)***, and cilia formation was evident by AT labeling (***Fig.4B***). Optical sectioning of the cilia was necessary to localize FPC inside the cilium. Notably, the antibody we used recognizes the FPC-CTD and this domain localized to the center of the cilia (***Fig.4C***, right). We also observed FPC in the nucleus, consistent with previous reports demonstrating that the FPC-CTD traffics to the nucleus (47). In comparison, *cpk* cells showed only minimum FPC staining in few cells, but not in the cilia (***Fig.4C***, bottom).

#### Loss of cystin affects ciliary architecture

Because both cystin and FPC are ciliary proteins, we examined how loss of cystin and FPC impacted cilia formation in *cpk* cells. For these experiments, we induced primary cilium development both in *w*t and *cpk* cells using serum starvation (39) and quantitated the fraction of cells with cilia (AT staining) relative to total cell number. We observed that the level of ciliation was very similar in *wt* and *cpk* cells with 77.35% of total *wt* and 75.82% of *cpk* cells exhibiting primary cilia ***(Fig.5A)***.

**Fig.5.**
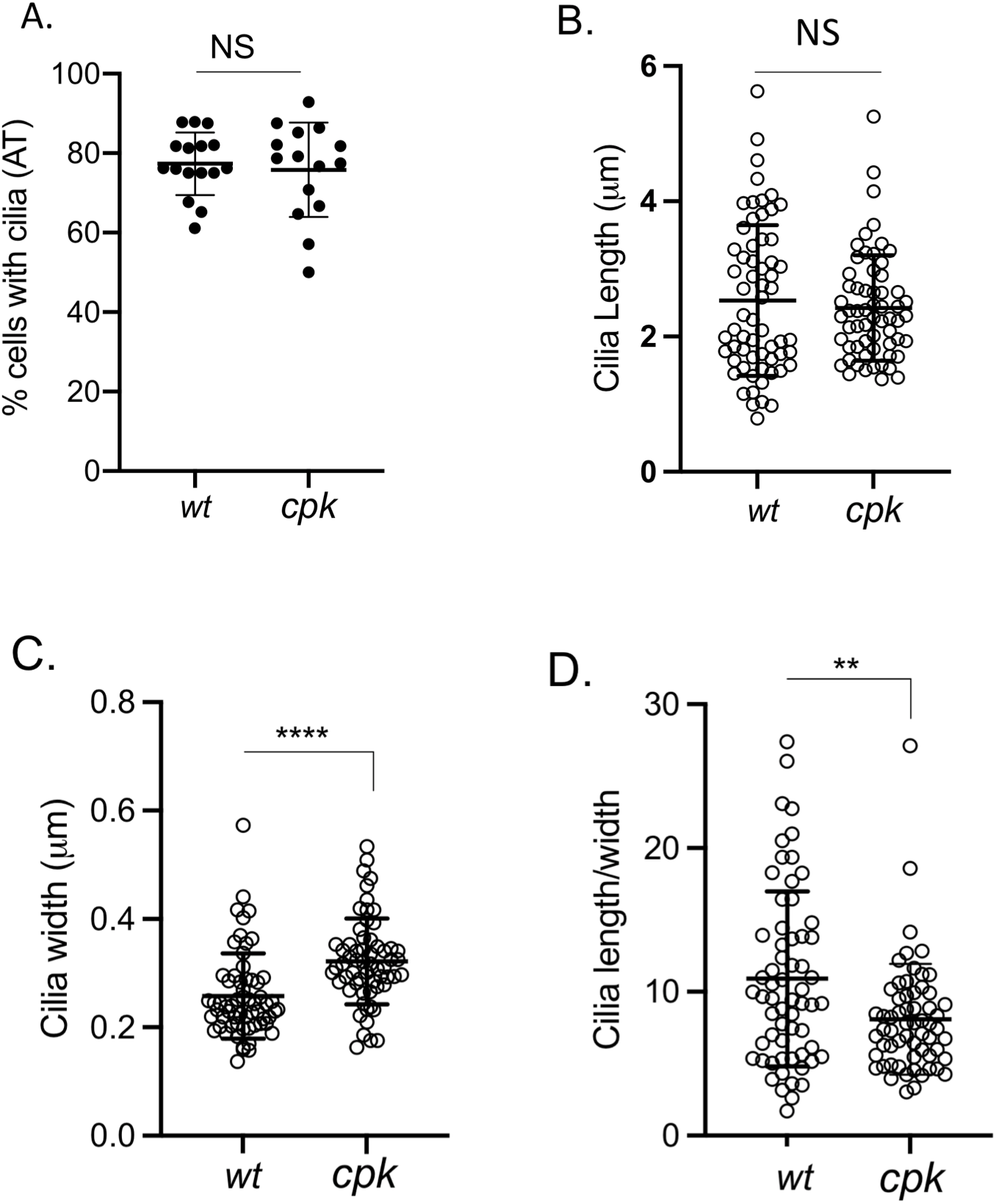
Loss of cystin in *cpk* cells leads to altered ciliary architecture. Acetylated tubulin (AT) staining was used to determine cilia number, thickness and length. **A:** Similar percentage of *wt* and *cpk* cells develop primary cilia. Cilia development was induced by serum starvation. Each data point represents the percentage of cells with cilia in a field with 22-44 cells in two experiments (*wt:* n=492, *cpk*: n=533). **B:** Cilia length**; C:** width (thickness, at the base of the cilia) were measured using ImageJ. Each data point represents measurements of one cilium (n=61, ****: p<0.0001;). **D:** Cilia length/width ratios are plotted from data points presented in B and C (n=61, **: p=0.0027).

Although there was no significant difference between the fraction of *wt* and *cpk* cells expressing cilia, the cilia of *cpk* cells structurally differed from those of *wt* cells ***(Fig.5 A-C)***. Morphometric analyses (length, width and width-to-length ratios) of cilia revealed no significant difference in the length of *wt* and *cpk* primary cilia ***(Fig.5B)***, but the *cpk* cilia were thicker at the base than *wt* (0.32 vs. 0.26 µm, p<0.0001) ***(Fig.5C)***, and the cilia length/width ratio was reduced in *cpk* cells (10.9 (*wt*), 8.1 (*cpk*), p=0.003) ***(Fig.5D)***. Therefore, the absence of cystin and significantly reduced FPC did not affect primary cilia formation, but was associated with altered ciliary architecture in *cpk* cells ***(Figs.4-5)***.

### II. The mechanism of FPC loss in *cpk* cells

#### Selective autophagy as a pathway for FPC removal

Relative *Pkhd1* mRNA levels were not lower in *cpk* kidneys and CCD cells compared to *wt* (***Fig.1-2)***. These results implied that FPC loss occurred at the protein level. If cystin plays an essential role in maintaining FPC levels, we reasoned that in the absence of cystin, FPC should undergo degradation through one of the intracellular protein degradation pathways (*i.e*. proteasomes, lysosomes, or autophagy).

Alternatively, enhanced FPC ectodomain shedding following notch-like processing could be a mechanism of FPC loss in *cpk* cells (47, 48). If FPC ectodomain shedding occurred in *cpk* cells, we would expect release of the FPC-CTD into the cytoplasm an/or its transport into nucleus. However, immunocytochemistry using antibodies specifically directed against the mouse FPC-CTD did not detect enhanced levels in the nuclei or the cytoplasm of *cpk* cells (***Fig.4A***). Neither did we observe increased FPC-CTD staining in the nucleus or cytoplasm when cells were treated with proteasome and lysosomes inhibitors. These results suggest that intracellular release of FPC-CTD was not increased.

Another possible mechanism by which cystin could modulate FPC levels is through vesicular trafficking by binding to the cytosolic side of FPC-containing vesicles. In this scenario, we would have expected colocalization of cystin and FPC, which was not observed (***Fig.4***). The absence of vesicle-associated cystin-FPC colocalization argues against cystin functioning as an escort protein of FPC-containing vesicles.

Loss of FPC might also occur during translation in the endoplasmic reticulum, e.g. endoplasmic reticulum-associated degradation (ERAD). However, proteasome inhibition with MG-132 for four hours did not increase levels of FPC in *cpk* cells, suggesting that FPC reduction in *cpk* cells does not involve ERAD ***(Fig.6A)***. However, increased levels of polyubiquitinated proteins in MG-132-treated samples of both *wt* and *cpk* cells indicated efficient proteasome inhibition, as shown later in ***Fig.8B***. Interestingly, prolonged (8 h) treatment with MG132 caused a severe reduction of FPC in both *cpk* and *wt* cells, but without affecting levels of cystoproteins Pc2 and Ift88 ***(Fig.6B,C)***. This prolonged MG132 effect is likely the result of selective autophagy activation as an alternative pathway for protein degradation (49-51). We found that inhibition of lysosomal fusion by chloroquine (CQ) did not alter the soluble fraction of FPC in either *wt* or *cpk* cells ***(Fig.6B)***.

**Fig.6.**
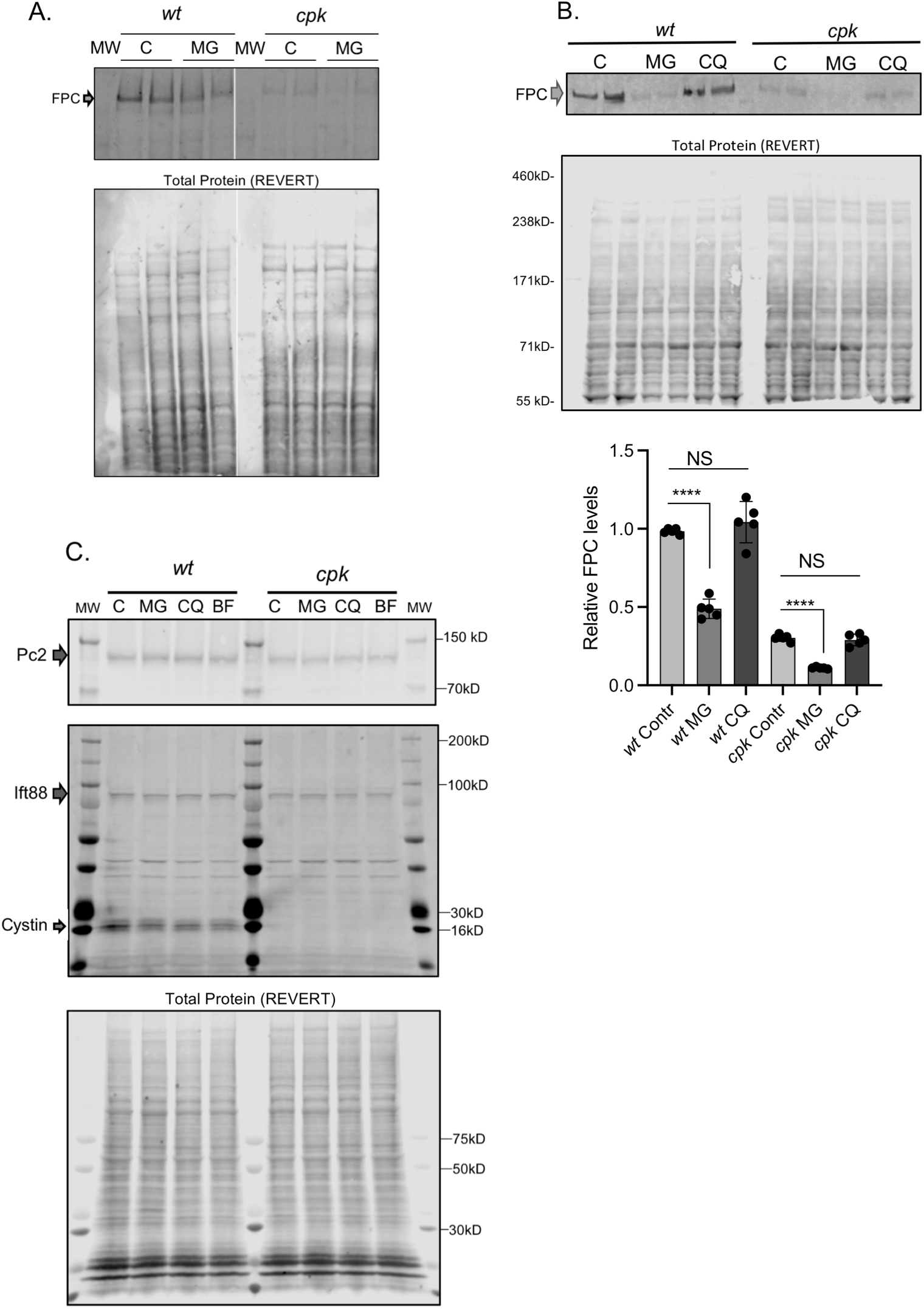
Proteasome or lysosome inhibition does not increase FPC levels. **A:** Representative gel is shown on top with biological duplicates for control (untreated) and proteasome inhibitor (MG, 4 hours) treated wt and *cpk* cells (top). Total protein stain (REVERT) shown to demonstrate equal loading. **B: Prolonged (8 hours) proteasome inhibition (MG) led to reduced FPC in *wt* and *cpk* cells**. Top shows a representative gel image (n=2). Middle shows a total protein gel image (Total Protein, REVERT). Quantitation of relative FPC levels/total protein shown at bottom. (N=3, n=2, ****: p<0.0001, unpaired *t* test). Lysosome inhibition (CQ) for 8 hours did not change FPC levels. **C: Levels of Pc2 and Ift88 did not change in the presence of proteasome (MG), or lysosome inhibitors (CQ, BF)**. MG, CQ and bafilomycin (BF) were applied for 4 hours, total protein (REVERT) staining is shown for reference (bottom).

To test if selective autophagy was activated following prolonged treatment with MG-132, we used SQSTM1/p62 as reporter. SQSTM1/p62 links the autophagy receptor LC3 with ubiquitinated proteins (52-54) to regulate cytoplasmic segregation and autophagic degradation of ubiquitinated proteins (55-57). In MG132-treated (8h) *wt* and *cpk* cells, we observed increased SQSTM1/p62 levels, consistent with activated autophagy. However, *cpk* cells appeared to have higher baseline levels of SQSTM1/p62, and also exhibited p62-oligomers, indicative of autophagy hyperactivation upon prolonged proteasome inhibition ***(Fig.7A)*** (58). Testing multiple samples of untreated *wt* and *cpk* lysates confirmed the higher abundance of SQSTM1/p62 in *cpk* compared to wt cells, suggestive of defective protein degradation and autophagy induction in cystin deficient *cpk* cells ***(Fig.7B)***.

**Fig.7.**
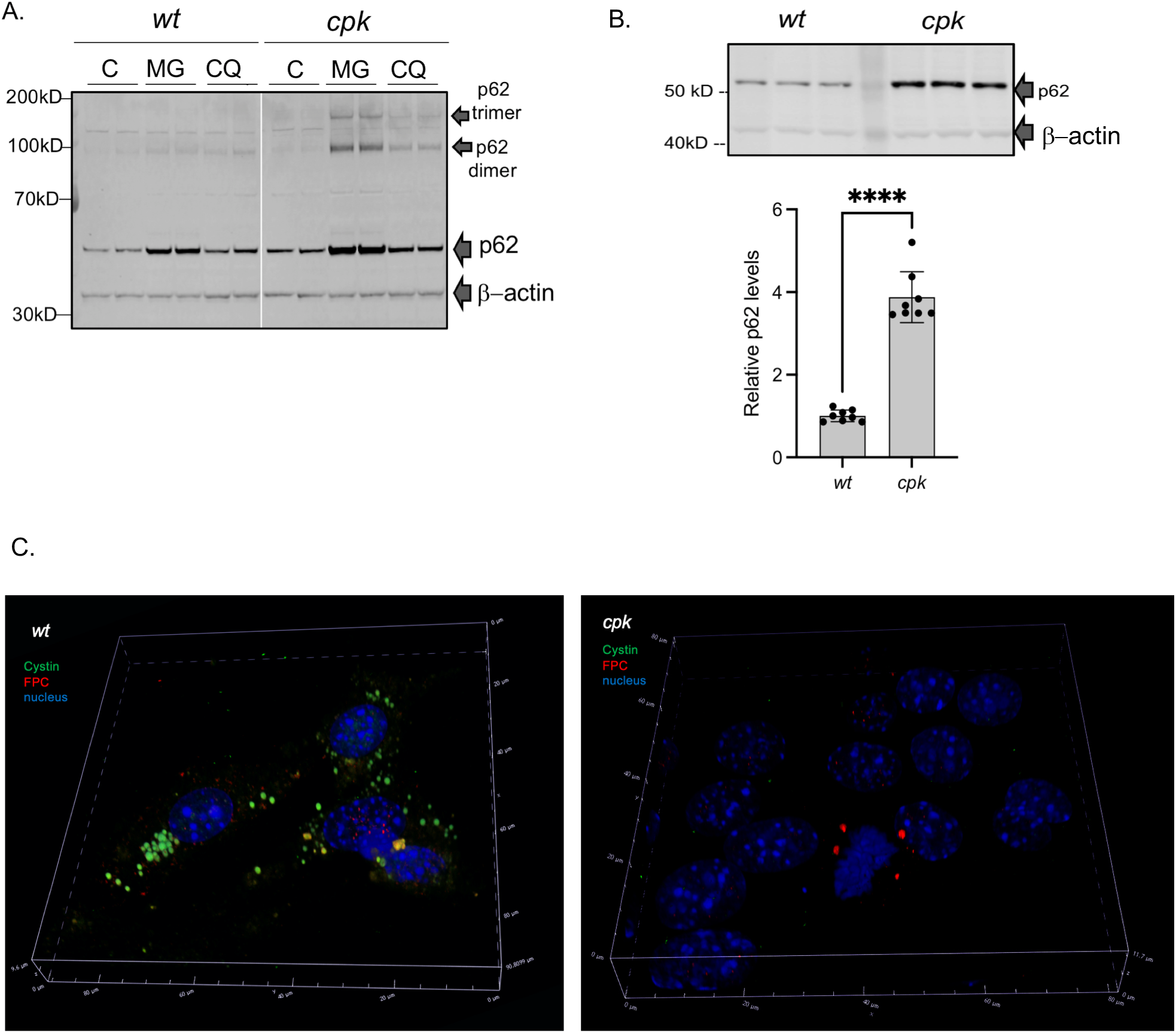
Autophagy as a pathway for FPC removal. **A: Hyperactivation of selective autophagy by proteasome inhibition, SQSTM1/p62 as reporter**. Representative gel image (n=2). C: control, MG: MG132, CQ: chloroquine. **B: Higher baseline SQSTM1/p62 levels in *cpk* compared to *wt* cells**. Representative gel image (top. Relative p62 levels/β-actin plotted on bottom (N=2, n=8), ****: p<0.000, unpaired *t* test). ***C:* Co-localization of cystin and FPC in cytoplasmic particles of *wt* cells following PI and LI**. *Wt* cells were stained with anti-cystin (green) and anti-FPC (red) antibodies following treatment with MG132 (8h) and CQ (4h), to facilitate formation of autophagic particles. Cystin colocalized with FPC in some particles (yellow staining). In cystin-negative *cpk* cells, smaller aggregates of FPC were detected in some cells (right).

**Fig.8.**
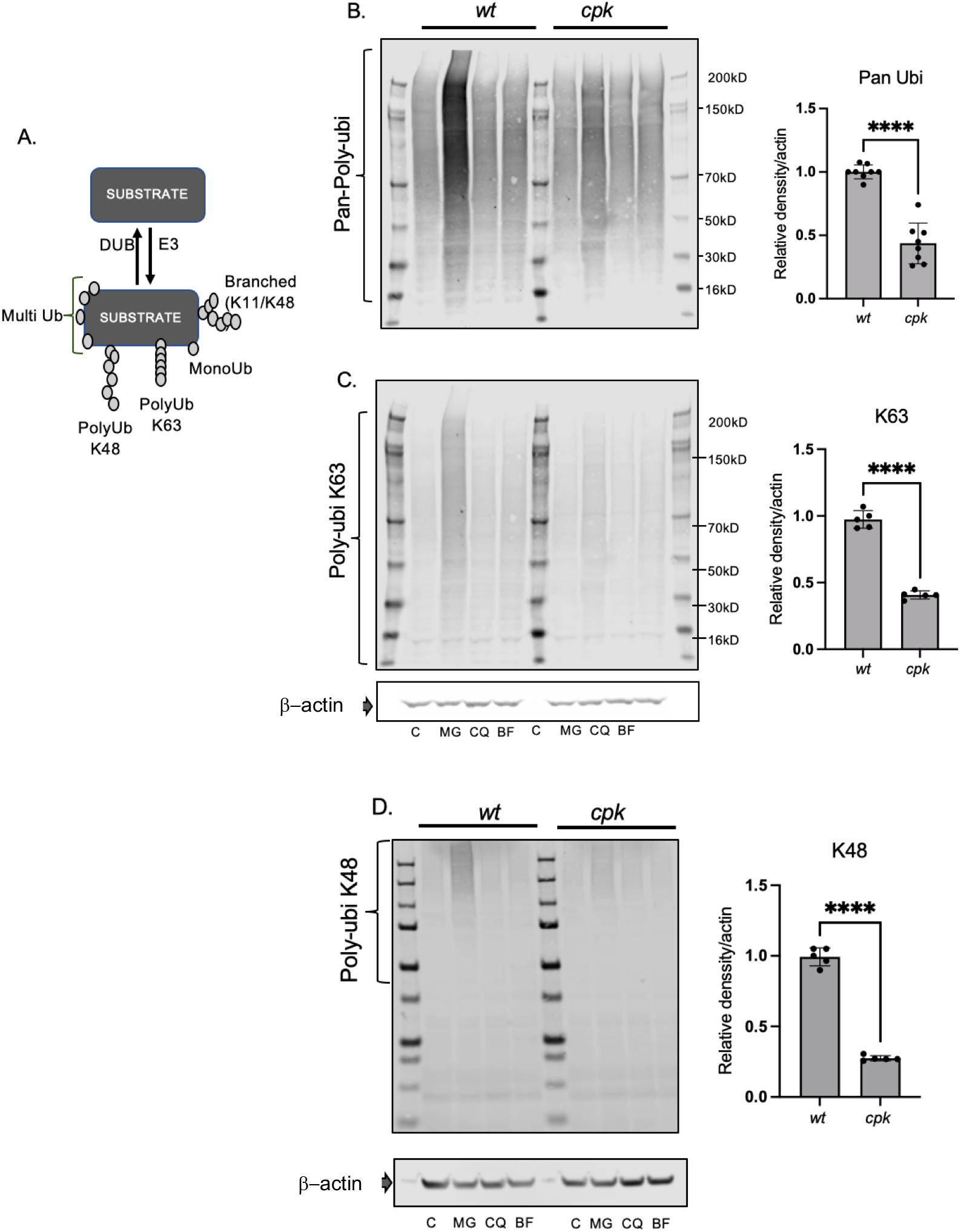
Reduced polyubiquitination in *cpk* cells. **A: Scheme depicting types of ubiquitination**. Proteins are polyubiquitinated prior to ERAD, lysosomal degradation or autophagy. The length of the poly-ubiquitin chain varies. **B-D: Reduced polyubiquitination in *cpk* cells**. Representative gels (left) are shown for polyubiquitination of cellular proteins in control (C), MG, CQ and BF-treated cells (4h). **B:** Pan-poly-ubiquitination (N=3, n=8); **C:** K63-linked (N=2, n=5); and **D:** K48-linked polyubiquitination (N=2, n=5). Results of densitometric analysis (right) are plotted relative to β-actin levels (****: p<0.00001, unpaired *t* test).

Because only three protein degradation systems exist in cells (proteasomal, lysosomal and autophagic, involving lysosomal fusion), we expected that when both the proteasomes and the lysosomes were inhibited, FPC would be sequestered in autophagic particles. To test this hypothesis, we treated the cells with MG-132 for 8 h to induce autophagy and then with CQ (4h) to inhibit lysosomal fusion, thereby preventing degradation of sequestered proteins before they fuse with lysosomes. We co-labeled cystin and FPC and observed large cystin-containing particles in *wt* cells, some of which also co-stained with FPC ***(Fig.7C, left)***, suggesting that both cystin and FPC were directed to autophagy. In *cpk* cells, we observed only few FPC-containing particles and no cystin ***(Fig.7C, right)***.

### III. The consequences of FPC loss in *cpk* cells

As described above, we have shown that cystin deficiency results in the loss of FPC (∼80%) and Pc2 (∼30%) and that re-expression of a GFP-tagged cystin can reduce cyst formation in r-*cpk* mice (4), but results only in partial recovery of FPC levels (***Fig.1)***. Therefore, we next sought to assess the functional consequences of FPC reduction in *cpk* cells. In previous collaborative work, we have shown that FPC is a functional component of the NEDD4-family of E3 ubiquitin ligase complexes, which facilitate the ubiquitination of multiple proteins, and that lack of FPC compromised the function of intracellular protein degradation systems (23). Therefore, ubiquitination may be compromised in *cpk* cells with minimal FPC levels, a condition which can itself lead to the activation of SQSTM1/p62-mediated selective autophagy (49). The data in the current study demonstrating higher SQSTM1/p62 levels in *cpk* cells compared to *wt* ***(Fig.7B***) are consistent with compromised protein degradation in *cpk* cells. Therefore, we compared the efficiency of polyubiquitination in *wt* and *cpk* cells.

#### Reduced ubiquitination in *cpk* cells

The NEDD4 family of E3 ligases belong to the <u>h</u>omologous to <u>E</u>6AP <u>C</u>-<u>t</u>erminus (HECT) domain E3 ligase group, which regulate multiple cellular activities through protein ubiquitination (for review: (59)). E3 ligases facilitate multiple types of ubiquitination (60) ***(Fig. 8A)***. FPC deficiency-associated E3 ligase dysfunction in *cpk* cells should, therefore, result in altered polyubiquitination of numerous proteins, impair the ubiquitin-proteasome system (UPS) as well as lysosomal/autophagosome degradative systems, leading to altered proteome homeostasis. We inhibited the proteasomes with MG132 and the lysosomes with CQ or BF for 4 hours and evaluated polyubiquitination, using linkage-specific anti-ubiquitin antibodies (61) to compare levels of polyubiquitination in *wt* and *cpk* cells. While proteasome inhibition led to increased ubiquitination in both wt and cpk cells, polyubiquitination was attenuated in *cpk* cells compared to *wt* (***Fig. 8B)***. Inhibition of the lysosomes with CQ or BF did not affect polyubiquitination and were excluded from further analysis.

To determine which type of ubiquitination was affected, we used anti-ubiquitin antibodies recognizing: 1) all types of polyubiquitin links (Pan-ubi), 2) K48-linked ubiquitin, which is the primary linkage type for proteasomal degradation, and 3) K63-linked ubiquitin, which is the primary linkage type degradation of cell surface receptors (62). *Cpk* cells exhibited lower levels of polyubiquitination in general ***(Fig. 8B)***, as well as reduced K63-linked ***(Fig. 8C)***, and K48-linked polyubiquitin chains ***(Fig.8D)***. These results are consistent with reduced E3 ligase activity in *cpk* cells, that may impede protein quality control by both the lysosomes and proteasomes and affect protein quality control at the endoplasmic reticulum and at the cell surface. Reduced polyubiquitination likely influences multiple cellular signaling pathways, as well as overall proteome homeostasis (59), making the consequences of cystin and FPC loss more complicated and severe than expected.

#### Enhanced expression of the epithelial sodium channel (ENaCα) in *cpk* cells

Proteomic analyses were beyond the scope of the current study. Therefore, as a proof of concept, we selected a functionally relevant NEDD4 E3 substrate, the epithelial sodium channel (ENaC) (24) to analyze the functional consequences of FPC loss in *cpk* cells. Of note, enhanced renal ENaC activity is associated with high blood pressure (63, 64), and systemic hypertension is a hallmark of ARPKD (25, 65). Previous studies have shown that expression of the functional alpha subunit of ENaC (ENaCα) is regulated via ubiquitination and degradation by both the UPS and lysosomes (30, 66, 67). Consistent with these results, we observed enhanced levels of ENaCα following both MG-132 and chloroquine (CQ) treatment for 4 hours ***(Fig. 9A)***. Interestingly, we also observed increased abundance of ENaCα in *cpk* compared to *wt* cells under control conditions ***(Fig. 9A,B)***. Confocal microscopy confirmed higher membrane ENaC levels in *cpk* cells ***(Fig.9C)***, with strong vesicular ENaCα staining and significant co-localization with lysosome-associated membrane protein 1 (LAMP1), which delivers proteins to the lysosome ***(Fig.9D)***. In *wt* cells, ENaCα was mostly observed in vesicular compartments, and was not colocalized with LAMP1. These data suggest that ENaC turnover is reduced in *cpk* compared to *wt* cells, consistent with previous findings indicating that FPC deficiency is linked to reduced NEDD4 E3 ligase activity, leading to defective degradation of cell surface transporters including ENaC (23).

**Fig.9.**
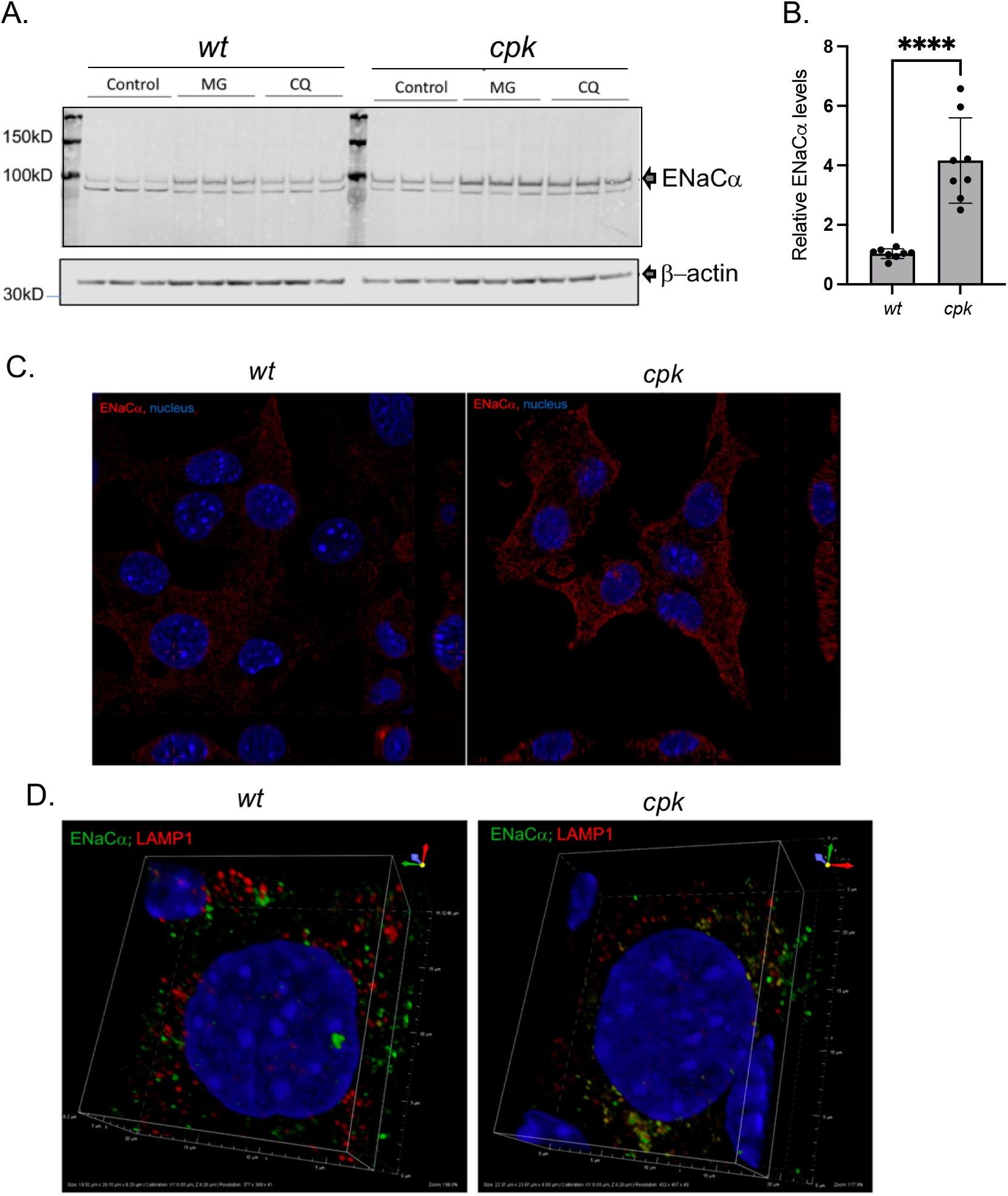
Higher ENaC levels in *cpk* cells. **A:** Representative WB of ENaCα in *wt* and *cpk* cells. β-actin was used as loading control. Both proteasome and lysosome inhibition (CQ) enhanced ENaCα levels, consistent with previously reported results. **B: Relative ENaCα levels**. (N=3, n=7; ***: p<0.0001). **C: Increased ENaC staining (red) in *cpk* cells**. Confocal microscopy images of *wt* (left) and *cpk* cells (right). **D: Co-localization of ENaC and LAMP1 (lysosomal marker)**. Minimal co-localization of ENaC (green) with LAMP1(red) in *wt* cells (left). ENaC (green) highly co-localized with LAMP1 (red) in *cpk* cells (right).

#### Higher ENaCα levels correspond to enhanced sodium transport

To determine whether increased ENaCα membrane abundance actually correlated with increased sodium transport, we performed whole-cell patch analysis on *wt* and *cpk* cells using previously published protocols (40, 41). Both cell types exhibited currents sensitive to the ENaC inhibitor amiloride ***(Fig. 10A-F)***, supporting the biochemical evidence for ENaC expression in both cell types ***(Fig.9)***. We measured net amiloride-sensitive currents by subtracting the currents in the presence of amiloride from control currents ***(Fig.10C,F)*** and plotted the mean I-V relation of amiloride-sensitive currents from *wt* and *cpk* cells, respectively ***(Fig.10G,H)***. Mean I-V curves of absolute amiloride-sensitive currents from *cpk* cells showed higher amiloride-sensitive currents at each membrane potential compared to *wt*. At positive membrane potentials (from -10 to 30 mV) when the I-V relation was expressed by current density (pA/pF, ***Fig.10H)***, we observed that *cpk* cells showed significantly larger amiloride sensitive currents, suggesting that there are more fluxes of amiloride-sensitive currents in *cpk* cells, representing more channels in the membrane. In addition, amiloride-sensitive currents from *cpk* cells are outward-rectified, in contrast to *wt* cells with inward rectified currents, contributing to high amiloride-sensitive currents from *cpk* cells at positive membrane potentials. Our findings indicate a positive correlation between higher ENaCα membrane abundance and enhanced amiloride-sensitive currents in *cpk* cells, a result consistent with the anticipated consequences of FPC deficiency-linked changes in proteome homeostasis.

**Fig.10.**
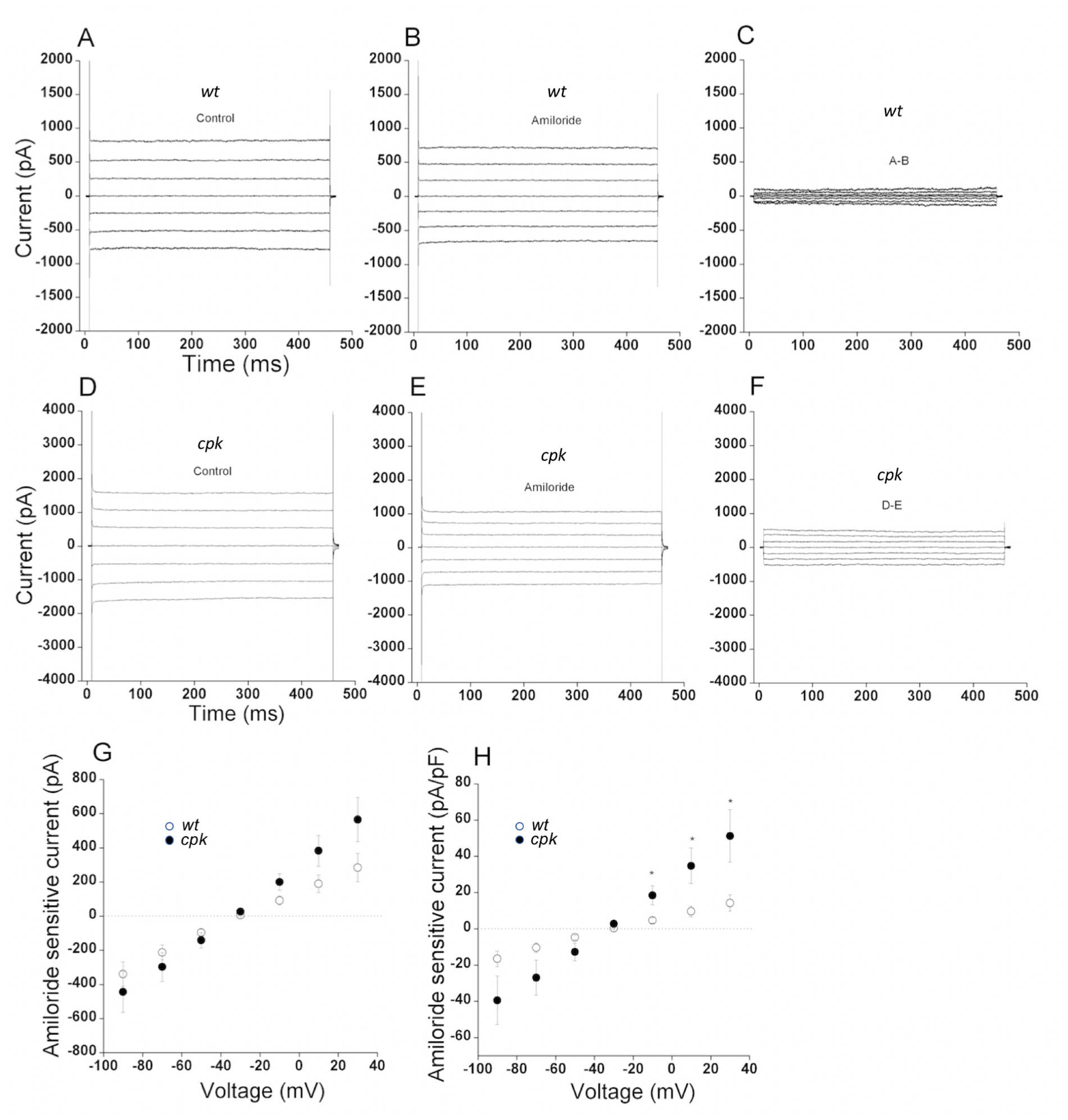
Higher relative amiloride sensitive (ENaC) currents in *cpk* compared to *wt* cells. Whole-cell currents were evoked by applying voltage steps from -90 to 30 mV in 20 mV increments. Cells were held at -30 mV. **A-C:** Representative whole-cell current recordings from *wt* cells; **D-F:** Representative whole-cell current recordings from *cpk* cells cells; **A and D:** Representative whole-cell current recordings in the absence of amiloride **B and E:** Representative whole-cell current recordings in the presence of amiloride (10 µM). **G:** Amiloride-sensitive currents (C and F) were measured by subtracting B from A and E from D. Mean I-V relationship of amiloride-sensitive currents expressed by total currents for *wt* (open circle, n=15) and *cpk* (dark circle, n=13) cells. **H:** Mean I-V relationship of amiloride-sensitive currents were expressed by current density for *wt* (open circle, n=15) and *cpk* (dark circle, n=13) cells. Values are means ±SE, *p<0.05.

## DISCUSSION

The current study demonstrates the cystin-dependent regulation of FPC levels in mouse kidneys and CCD cell lines, providing the first evidence of a mechanistic connection between the phenotypes of *cpk* mice and patients with ARPKD caused by *PKHD1* as well as *CYS1* mutations. Our studies demonstrate that in mouse kidneys and CCD cell lines, loss of cystin led to an 80-90% reduction in FPC levels, which occurred at the protein level. Both the natural mutation-mediated loss of cystin in *cpk* kidneys and CCD cells, as well as siRNA removal of *Cys1* mRNA in *wt* CCD cells led to FPC reduction. These results (*in vivo, in vitro in cpk* and siRNA of *Cys1* in *wt* cells) argue against the possibility that reduced FPC in *cpk* cells is the result of a *cpk* CCD cell line-specific phenomenon. Notably, collecting duct-specific expression of the cystin-GFP fusion protein in r-*cpk* mice (4) only partially rescued FPC levels ***(Fig.1A)***. The partial, but statistically highly significant (p<0.0005) recovery of FPC levels in whole kidney lysates from *r-cpk* mice is consistent with the collecting duct-restricted expression of the cystin-GFP construct (4). It is also possible that, despite its high level of expression, the cystin-GFP fusion protein does not have comparable function as endogenous cystin, as has been demonstrated for multiple GFP-tagged proteins with reduced, altered, toxic or dominant negative effects (42, 43, 68, 69). However, our observation that depleting cystin through siRNA in *wt* cells, which express endogenous cystin and FPC, caused a cystin-depletion-dependent FPC reduction, support the thesis that cystin is necessary to stabilize FPC and when cystin is removed, FPC levels quickly decrease. These data and the observation that FPC-deficiency did not reduce cystin expression in any of the tested *Pkhd1* mutant mouse kidneys suggest that cystin-mediated FPC expression regulation is unidirectional.

To our knowledge, no prior studies have analyzed primary cilia development in *cpk* cells. We demonstrate that loss of cystin and FPC did not disrupt ciliogenesis, but the structural architecture of the cilia was different in *cpk* vs. *wt* cells ***(Fig.5***). Further investigation will be required to define the mechanistic alterations that underpin this novel observation and whether ciliary structural alterations contribute to the *cpk* renal cystic phenotype.

In order to understand the mechanisms of FPC loss in the absence of cystin, we examined the role of intracellular protein degradation systems. We could not recover FPC by inhibiting the proteasomes or lysosomes in *cpk* cells ***(Fig.6)***. These data implicate autophagy as the only remaining mechanism responsible for FPC removal in the absence of cystin. However, the very low baseline levels of FPC in *cpk* cells confounded our ability to experimentally analyze this mechanism in cystin-deficient cells. Moreover, we note that if FPC is a functional component of the cellular protein quality control, as proposed by Kaimori *et al* (23) and supported by our results (***Figs 8-10***), it may not be possible to recover FPC by simply inhibiting individual protein degradation systems, or with simultaneous pan-inhibition, because these systems are functionally linked at multiple levels and their integration is required for cellular homeostasis [for reviews: (70-72)]. Inhibition of one protein degradation pathway activates alternative degradation pathways (49) and stress responses, leading to alterations in cellular functions, including translation inhibition (73-75). In addition, once a protein is sequestered in autophagic particles, it may not be resolved by standard cell lysis conditions (76). Analysis of protein aggregates would require FPC domain-specific antibodies and those are not available for endogenously expressed FPC. Therefore, it was not feasible to further investigate the nature and localization of such sequestered complexes at this time.

We observed that SQSTM1/p62-regulated selective autophagy was activated in cystin-deficient *cpk* cells at baseline ***(Fig.7B)***, most likely by disturbances in E3 ligase function ***(Fig.8)***, which is necessary for proper protein quality control [for review: (59, 71)]. The formation of protein aggregates when we hyperactivated autophagy and inhibited the lysosomal fusion of the particles ***(Fig.7C)***] also suggests that autophagy is responsible for FPC removal. Moreover, we found that autophagy hyperactivation led to the degradation of FPC not only in *cpk*, but also in *wt* cells ***(Fig.6B)***. Our results are also consistent with previous reports indicating that degradation of ciliary proteins occurs through selective autophagy (77, 78). Further studies will be necessary to define the functional mechanisms by which cystin shelters FPC from degradation and contributes to the maintenance of the cellular proteome. Although we do not understand all facets of the cystin-FPC interaction, the current studies, performed in mouse kidneys and CCD cell lines expressing endogenous FPC, clearly demonstrate that loss of cystin leads to significant FPC reduction with severe consequences. Conversely, FPC loss does not alter cystin expression in *Pkhd1* mutant mice (***Fig. 1D***) which lack classic ARPKD phenotype (6, 7).

As an important consequence of cystin and FPC loss, we demonstrated significant reduction in polyubiquitination **(Fig.8)**, supporting the findings of Kaimori and colleagues (23) that FPC plays crucial role in protein quality control and proteome management. Our data are also consistent with the interactive feedback among intracellular protein degradation systems (proteasome, lysosomes and different types of autophagy) (79) in maintaining proteostasis (80). When one of these systems is rendered dysfunctional, other protein degradative pathways undergo dynamic adjustments to maintain proteome integrity (71). We propose that in the absence of cystin, FPC undergoes targeted degradation that renders the NEDD4 family of E3 ligase complexes dysfunctional. The downstream consequences involve dismantling of the integrated protein quality control mechanisms that maintain cellular proteome integrity. This hypothesis is supported by previous studies reporting attenuated expression of epithelial adhesion molecules (81, 82)}, altered basement membrane deposition (83), and metabolic abnormalities (8) in *cpk* cells.

Proteomic analysis, combined with genomics, will be necessary to fully elucidate the extent and severity of consequences caused by cystin deficiency and the consequent reduction of FPC deficiency on the cellular proteome. Such studies will not only improve our understanding of the complexity of mechanisms leading to cyst formation, but will also provide important new insights about the function of FPC and the FPC-containing E3 ligase complexes. As an initial proof of concept, we examined a functionally relevant NEDD4 E3 substrate, the epithelial sodium channel (ENaC) (24). We observed elevated ENaC protein levels and increased epithelial sodium transport in *cpk* cells, which supports our hypothesis and reveals a possible mechanism underlying the systemic hypertension that is a hallmark symptom of ARPKD (25, 65).

Our studies support the proposition that cystin maintains FPC levels, and FPC is required for the physiological function of certain HECT-domain E3 ubiquitin ligase complexes, especially in renal epithelial cells. We have extended the previously proposed role of FPC in maintaining the integrity of signaling pathways, protein quality control, and turnover (23), now to include cystin. We have shown that cystin affects FPC abundance in a unidirectional manner and that loss of FPC in *Pkhd1* mutant mouse kidneys was not associated with reduced cystin levels. This observation may be critically important for clarifying an otherwise apparent paradox. That is, why does the loss of FPC cause renal cysts in *cpk* but not *Pkhd1* mutant mice? We propose that, in mice, cystin has at least two essential roles: maintenance of FPC as a required functional component of the NEDD4 E3 ligase complexes, and inhibition of *Myc* expression via interaction with necdin, as we have demonstrated previously (18). In mouse kidneys, more specifically in CCD epithelial cells, both FPC loss and Myc overexpression are required for cystogenesis. By contrast, *Pkhd1* mutant mice lacking FPC retain normal cystin levels (**Fig.1**), which can keep *Myc* expression low through the cystin-necdin interaction (18, 31). Our results imply that disruption of proteome homeostasis is a significant pathophysiological consequence of FPC loss, yet not sufficient for renal cyst formation in mice when cystin is present. Identification of all E3 ubiquitin ligases, components of the complexes, and the mechanisms by which the cellular proteome is affected by the loss of FPC in mouse and human kidneys will require further investigations. Those studies will likely shed light on factors that are responsible for the differences observed between human and mouse kidney phenotypes following FPC loss.

Finally, our data establishing a functional link between cystin and FPC expression in mouse kidneys and CCD cells, as well as the recent identification of patients with *CYS1*-related ARPKD, support the use of the *cpk* mouse as an important experimental model to study ARPKD pathogenesis and experimental therapeutics. Our studies also provide novel perspectives on the renal phenotype in mouse models of ARPKD, suggesting that future mechanistic studies should evaluate the interactome that underpins FPC and cystin expression, *Myc* transcriptional regulation, and maintenance of the cellular proteome, as well as how disruption of these interacting pathways drives ARPKD pathogenesis.

## AUTHOR CONTRIBUTIONS

**Yiming Zhang** contributed to experimental design, performed most of the experimental procedures, contributed to data analysis and manuscript writing. **Chaozhe Yang** designed experiments, prepared samples, and performed western blotting analysis of cystin and FPC expression in mouse kidneys. **Wei Wang** designed, performed, and analyzed the patch clamp studies. **Naoe Harafuji** designed the experiments and performed and analyzed mRNA expression levels in cells and kidney tissues. **P. Darwin Bell** developed the immortalized cell lines and helped to interpret initial results. **E. Sztul’s laboratory (P. Stasiak) performed the cilia morphology studies. Lisa M. Guay-Woodford** supervised studies in *wt* and *cpk* mice, contributed to the interpretation of results and manuscript writing. **Zsuzsanna Bebok** conceived the study, co-designed the majority of the experiments, supervised the analysis and presentation of experimental results, and wrote the manuscript.

## Acknowledgments

We thank Dr. Adam Richman for critical reading and help with editing of the manuscript.

## Funding

UAB HRFD Core Center and O’Brien Center for Acute Kidney Injury Center Pilot Program (ZsB) P30DK072482, UAB CF Research and Translation Core Center and CCF-ROWE19RO, UAB Research and Development Program (ZsB and WW); R01GM122802 (ES). R01DK121530, PKD Foundation, and The Moran Family Foundation (LGW).

## Notes

### Competing Interest Statement

The authors have declared no competing interest.

### Summary of Updates

We provide novel data on the role of cystic and FPC expression on primary cilia development (Fig.5). The text has been revised to provide details about experimental conditions.

